# FGF21 signals through KLB-expressing glutamatergic neurons in the hindbrain to mediate the effects of dietary protein restriction

**DOI:** 10.1101/2025.04.19.649640

**Authors:** Redin A. Spann, Sora Q. Kim, Md Shahjalal H. Khan, Diana A. Albarado, Sun O. Fernandez-Kim, Hans-Rudolf Berthoud, David H. McDougal, Heike Münzberg, Yanlin He, Sangho Yu, Christopher D. Morrison

## Abstract

Animals adaptively respond to protein restriction by altering both behavior and metabolism. Previous work demonstrated that the metabolic hormone FGF21 acted in the brain to coordinate these adaptive responses, but the exact site of action remains unclear. Here, we identify a discrete population of glutamatergic, *Klb*-expressing neurons in the nucleus of the solitary tract (NTS), demonstrating that these neurons are key to mediating FGF21 action during protein restriction. Using a novel *Klb-Flp* mouse line combined with intersectional genetics, we demonstrate that these neurons are directly activated by FGF21. While previous work implicated the SCN, PVN, and VMH in FGF21 action, we find that these areas do not impact the response to protein restriction. Instead, selective ablation of NTS-KLB neurons prevents metabolic adaptations to protein restriction (food intake, food choice, and energy expenditure), while their chemogenetic activation is sufficient to drive these responses. These findings establish NTS-KLB neurons as a critical node for detecting protein status and coordinating whole-body metabolic responses, providing new insight into how the brain monitors and maintains protein homeostasis.

## Introduction

Fibroblast growth factor 21 (FGF21) is a liver-derived endocrine hormone that influences energy expenditure, food choice and metabolism. We have previously shown that FGF21 plays a key role in the detection of the protein restricted state ^1, 2, 3^. Protein restriction triggers a variety of adaptive physiological responses, including changes in growth, energy expenditure, food intake, food choice, and motivation for protein. These responses act to protect the animal from the protein restricted state, but they also improve metabolic health and enhance longevity in rodents ^4, 5^. Importantly, FGF21 is critical for these adaptive responses to protein restriction, such that the effects of protein restriction on food choice, motivation to consume protein, energy expenditure, and even lifespan are lost in mice lacking FGF21. Collectively, these data strongly suggest that FGF21 is a signal of the protein restricted state that serves to coordinate a variety of adaptive responses to protein restriction.

FGF21 signals through a receptor system consisting of the FGF receptor 1c (FGFR1c) and the co-receptor beta-klotho (KLB). While FGFR1c mediates intracellular signaling, KLB mediates extracellular binding and thus provides cellular specificity for FGF21 signaling ^6, 7, 8^. FGFR1c and KLB are co-expressed in only a few tissues, including adipose tissue, pancreas, lung, and brain ^9, 10^. Importantly, our previous work indicates that *Camk2a-Cre* mediated deletion of KLB from neurons blocked the effects of dietary protein restriction on energy expenditure, glucose handling, and protein preference and motivation ^2^. These data strongly suggest that the brain is a primary mediator of FGF21 action in the context of dietary protein restriction, consistent with prior work indicating that the brain mediates the effects of pharmacological FGF21 administration ^10, 11, 12, 13^.

Although it is well accepted that the brain plays a critical role in mediating FGF21 action, the specific brain areas mediating FGF21 signaling have remained unclear. Analysis of *Klb* expression suggests that *Klb* is robustly expressed in two brain areas, the suprachiasmatic nucleus (SCN) and the nucleus of the solitary tract (NTS) ^9, 10^, although lower levels of *Klb* expression are detected in several additional locations both within and outside the hypothalamus. Importantly, early work suggested that the NTS was not critical for mediating FGF21 action ^10^, and thus recent attention has focused on other brain areas, including the SCN, PVN, and VMH ^10, 12, 14, 15, 16, 17, 18^.

Here, we describe our strategy, using the physiological model of dietary protein restriction, to identify the brain areas that mediate FGF21 action. Our data demonstrate that FGF21 signaling in the PVN or VMH is dispensable, but that a discrete population of glutamatergic *Klb*-expressing neurons in the NTS is required for the response to protein restriction. Deletion of these NTS-KLB neurons blocks changes in food intake, food choice, and energy expenditure in response to a low protein diet, while their chemogenetic activation increases both food intake and energy expenditure. These data demonstrate that glutamatergic NTS-KLB neurons are responsive to FGF21 and critical for adaptive responses to dietary protein restriction.

## Results

### *Klb-Flp* drives recombination within two distinct GABAergic and Glutamatergic neural populations

To determine where FGF21 signals within the brain, we generated a novel mouse line in which Flp recombinase is inserted into the 3’ end of the beta-Klotho *Klb* gene, connected with the T2A linker (**Figure 1 A**). This genetic manipulation results in FLP production wherever *Klb* is expressed. These *Klb-Flp* mice were then crossed with the dual recombinase *RC::FLTG* (*FLTG*) reporter line ^19^. This intersectional approach provides additional power for phenotyping KLB neurons since Flp-based recombination of FLTG drives the expression of tdTomato, but the addition of Cre removes tdTomato while activating GFP expression ^19^.

**Figure 1.**
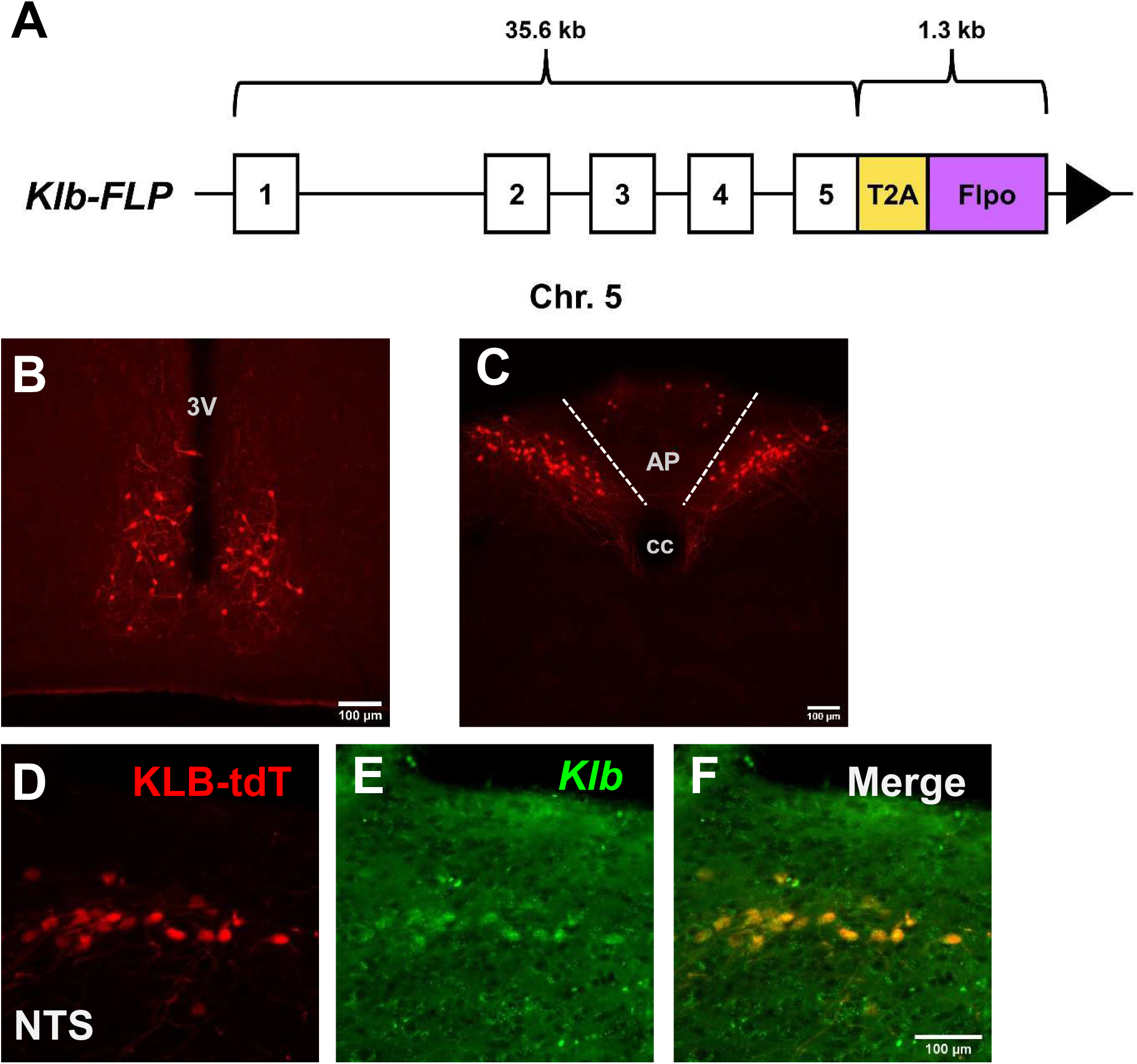
*Klb-Flp* mice mark two neural populations in the brain. A novel mouse was created that expresses Flp recombinase driven by *Klb* expression (A). *Klb-Flp* mice were then crossed with FLTG reporter mice and two populations of tdTomato-positive neurons were observed: in the suprachiasmatic nucleus of the hypothalamus (SCN) (B) and the nucleus of the solitary tract (NTS) (C). *Klb* expression was confirmed using RNAscope reagents in reporter mice (D-F).

Assessing Flp-induced recombination (tdTomato expression) within the brain of *Klb-Flp;FLTG* mice revealed prominent tdTomato expression within only two areas of the brain: the suprachiasmatic nucleus (SCN) and the nucleus of the solitary tract (NTS (**Figure 1 B&C)**. Beyond the brain, *Klb-Flp;FLTG* mice reveal tdTomato expression in other tissue known to express Klb, including the liver (**Figure S1**). To confirm that these *Klb-Flp* neurons actively express *Klb*, we used RNA-scope to colocalize *Klb* mRNA with tdTomato expression in *Klb-Flp;FLTG* mice (**Figure 1 D-F**). Within the NTS, we observe essentially 100% colocalization between these markers (**Figure 1 F**), indicating that *Klb-Flp* neurons within the NTS actively express *Klb*.

Intersectional genetics was then used to further phenotype *Klb-Flp* neurons by crossing *Klb*-*Flp*;FLTG mice with various Cre recombinase mouse lines. Since the adaptive response to protein restriction depends on Klb expression in *Camk2a*-positive cells ^2^, we first tested whether *Camk2a-Cre* would induce recombination (removal of tdTomato and expression of GFP) in *Klb-Flp* cells. We found that both the SCN and NTS populations expressed GFP but not tdTomato (**Figure 2 A&B**). *Klb-Flp;FLTG* mice were then crossed with established *Vglut2-Cre* (glutamatergic) and *Vgat-Cre* (GABAergic) lines. Interestingly, *Vglut2-Cre* induced recombination (GFP expression) only within the NTS (**Figure 2 C&D)**, while *Vgat-Cre* induced GFP expression only within the SCN (**Figure 2 E&F**). These intersectional crosses suggest that both SCN and NTS *Klb-Flp* neurons express *Camk2a-Cre*, but that the NTS population is glutamatergic while the SCN population is GABAergic.

**Figure 2.**
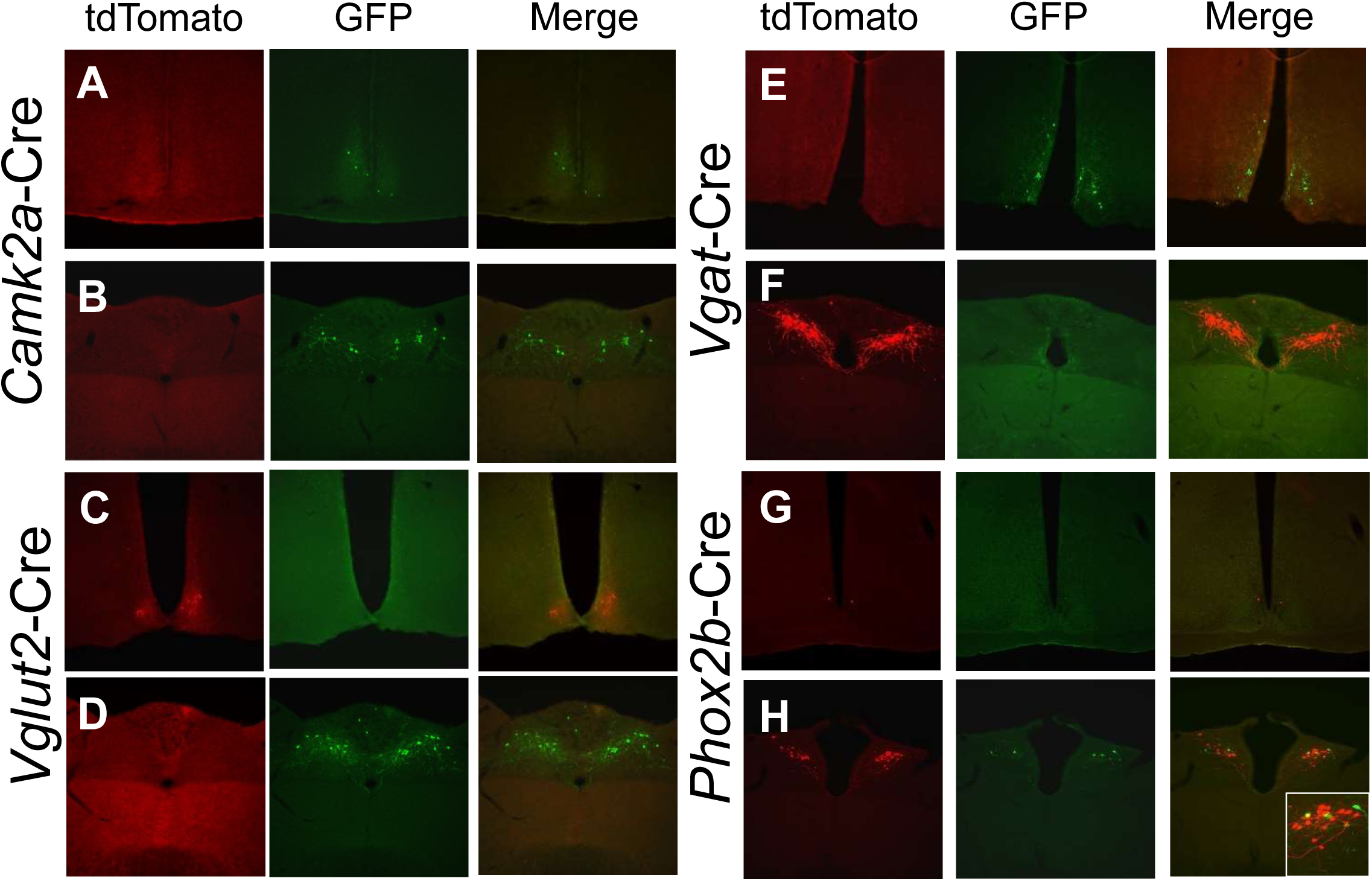
Characterizing KLB neurons with a dual recombinase approach. *Klb-FLP;FLTG* mice were crossed with various Cre mouse lines to further characterize Klb neurons via intersectional genetics. Crossing with *Cam2ka-Cre* mice resulted in GFP expression in the SCN (A) and NTS (B), indicating co-expression of *Klb* and *Camk2a.* Crossing with *Vglut2-Cre* mice resulted in GFP expression in the NTS but not the SCN (C&D), indicating that NTS-KLB neurons are glutamatergic. Crossing with *Vgat-Cre* resulted in GFP expression in the SCN but not NTS (E&F), indicating that SCN-KLB neurons are GABAergic. Lastly, *Phox2b-Cre* induced GFP expression in some but not all NTS-KLB neurons (G & H).

The *Phox2b-Cre* has been previously used to target the dorsal vagal complex ^20^, and we therefore tested whether *Phox2b-Cre* would selectively target *Klb-Flp* neurons in the NTS. As expected, *Phox2b-Cre* based recombination (GFP-induction) was not detected in the *Klb-Flp* neurons in the SCN (**Figure 2 G)**. However, *Phox2b-Cre* also only targeted some of the *Klb-Flp* neurons within the NTS (**Figure 2 H)**. Thus *Phox2b-Cre* is not an effective tool for targeting the entirety of NTS FGF21 signaling in this *Klb-Flp* mouse line.

### KLB neurons in the NTS are responsive to FGF21

Since the above data highlight the NTS and SCN as likely locations for FGF21 signaling, we next tested whether FGF21 impacts the activity of these neuronal populations. IP injection of FGF21 (1mg/kg) had no effect on cFos induction within the SCN (**Figure 3 A, B, & F**), but robustly induced cFos expression within nearly all *Klb-Flp* neurons in the NTS (**Figure 3 C, D**, **& F**).

**Figure 3.**
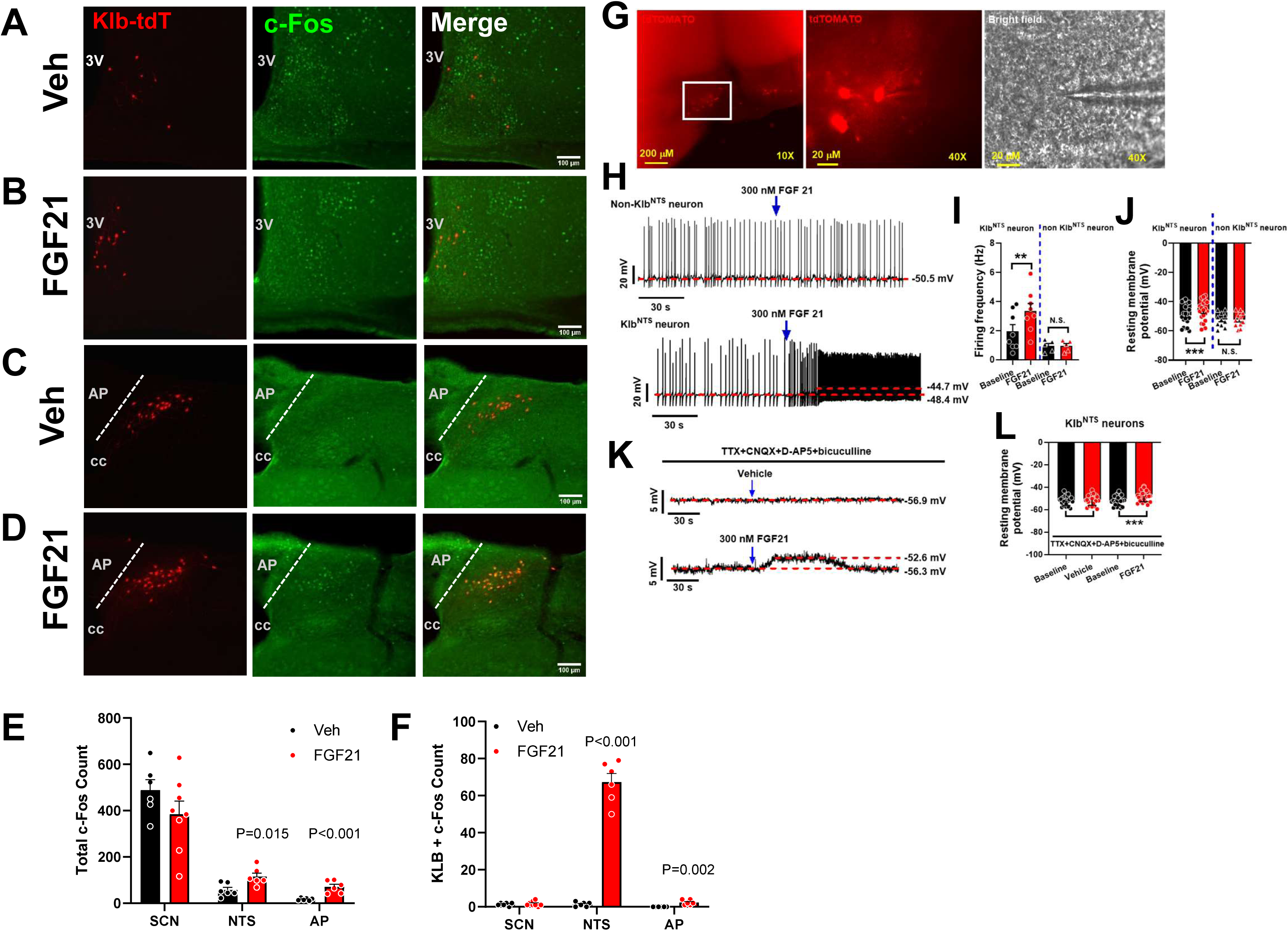
NTS-KLB neurons are responsive to FGF21. *Klb-Flp;FLTG* mice were given an i.p. injection of hFGF21 or vehicle, and IHC was performed in the brain to colocalize tdTomato expression with c-Fos as a marker of neural excitation. In the SCN, there was no change in total c-Fos or c-Fos colocalization with tdTomato (A, B, E & F). In the NTS and AP, however, FGF21 injection increased total c-Fos and the coexpression of c-Fos and tdTomato (C, D, E & F). Slice patch-clamp electrophysiology was also performed in the NTS of reporter mice with administration of hFGF21 (300nM, G). FGF21 increased both firing rate and membrane potential in tdTomato-positive neurons compared to baseline (H, I, & J). KLB neurons were still activated when preincubated with a cocktail of inhibitors (K & L).

Electrophysiology was then used to directly record from *Klb-Flp* neurons in ex vivo slices. Within the NTS, FGF21 (300nM) produced a robust increase in firing frequently in KLB neurons, but not nearby unlabeled neurons (**Figure 3 H & I**). This increase in firing frequency was associated with an increase in resting membrane potential that was insensitive to inhibitors of synaptic signaling (**Figure 3 K & L**). Similar work in the SCN produced more divergent responses, with FGF21 increasing the firing rate in some *Klb-Flp* SCN neurons, but either reducing or having no effect on firing in others (**Figure S2**). Together, these data indicate that FGF21 can directly activate *Klb-Flp* neurons within the NTS but produces inconsistent effects within the SCN.

### Glutamatergic KLB neurons mediate responses to protein restriction

Since *Klb-Flp* neurons in the NTS and SCN are differentially marked by *Vglut2-Cre* and *Vgat-Cre* (Fig 2), these Cre lines were used to selectively delete Klb to determine the impact on the adaptive response to protein restriction. Male *Vglut2-Cre*; *Klb^lox/lox^* (*Klb^Vglut^*) mice and Cre negative littermate controls (*Klb^lox/lox^*) were placed on a control diet (Con, 20% protein) or isocaloric low protein diet (LP, 5% protein) at 8 weeks of age, and remained on this diet for 5 weeks. We first confirmed that Klb expression was reduced in the NTS with qPCR (P=0.0021, **Figure S3 A**). Next, we also measured mRNA in the liver to confirm that mice responded to LP diet. As expected, expression of *Fgf21* was increased in both *Klb^lox/lox^* and *Klb^Vglut^* animals along with phosphoglycerate dehydrogenase (*Phgdh)* and asparagine synthetase (*Asns)* indicating stress in amino acid metabolism (**Figure S3 B**). In male *Klb^lox/lox^*mice, LP diet reduced body weight gain and lean mass gain (P < 0.05, **Figure 4A & D**), while also increasing total food intake (P < 0.01, **Figure 4 B**). These LP-induced effects were completely lost in *Klb^Vglut^* mice.

**Figure 4.**
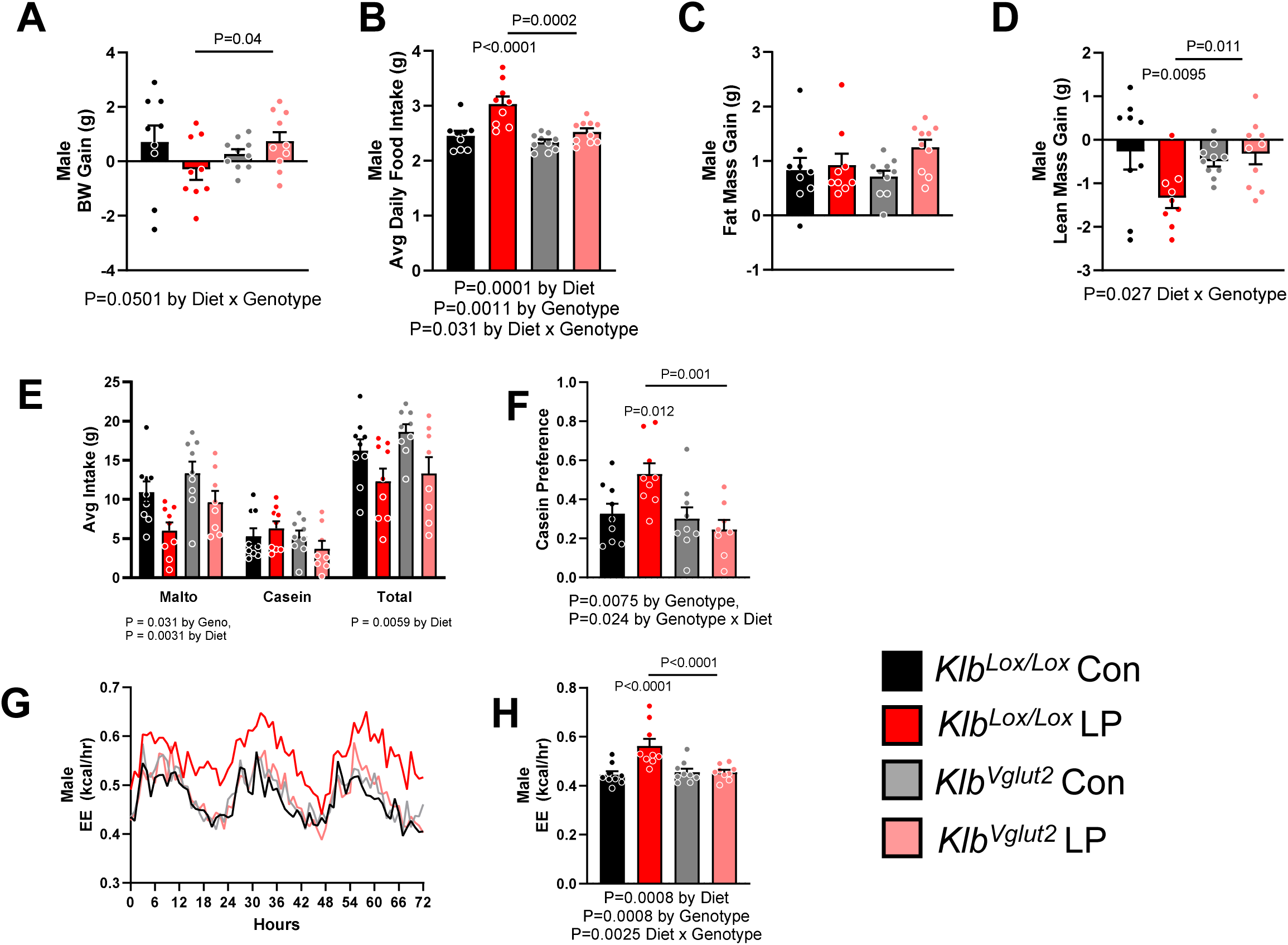
Glutamatergic KLB-positive neurons mediate responses to protein restriction. In male mice, *Vglut2-Cre* specific deletion of *Klb* (*Klb^Vglut2^*) blocked the response to low protein diet. *Klb^Vglut2^* mice did not lose weight when faced with dietary protein restriction compared to control littermates (*Klb^lox/lox^*, A). Specifically, this disparity was mainly due to lean mass (D) and not fat mass (C). *Klb^Vglut2^* mice also did not respond to low protein diet by consuming more food (B) and did not have an increased preference for protein (casein) (E & F) compared to wild-type mice. Lastly, male *Klb^Vglut2^* mice did not have an increase in energy expenditure (EE) when on low protein diet compared to wild-type mice (G & H). All data are mean ± SEM, analyzed with two-way ANOVA and post-hoc T-test, Avg EE was analyzed by ANCOVA and post-hoc T-test. n=8-10/group.

Protein restriction also alters macronutrient preference by increasing the preference/consumption of high-protein foods ^1, 2, 21^. Macronutrient preference was tested in male mice by giving them simultaneous access to sippers containing either 4% casein (protein) or 4% maltodextrin (carbohydrate). In *Klb^lox/lox^*mice, LP decreased maltodextrin consumption while increasing casein consumption (**Figure 4 E**), significantly increasing preference for casein (P = 0.012, **Figure 4 F**). Again, this LP-induced increase in casein preference was absent in *Klb^Vglut^* mice. (**Figure 4 E & F**). Finally, total energy expenditure (EE) was measured via indirect calorimetry. As previously observed, LP diet significantly increased EE (BW adjusted via ANCOVA) in *Klb^lox/lox^* mice (P< 0.01, **Figure 4 G & H**), but this effect was also lost in *Klb^Vglut^* mice. These data indicate that LP-induced changes in growth, food intake, macronutrient preference, and energy expenditure all require *Klb* expression within glutamatergic neurons, consistent with previous work ^22^.

Prior work indicates that protein restriction does not impact the same array of endpoints in females as in males ^23, 24, 25^. Here, the LP diet did not affect BW gain but did increase food intake in female *Klb^lox/lox^* mice (P < 0.01, **Figure S4 A-D**). However, this LP-induced hyperphagia was lost in female *Klb^Vglut^* mice. Therefore, while the response to LP diet may be different in females, these effects nevertheless require KLB signaling in glutamatergic neurons. Despite these differences in behavioral responses to low protein, we did not detect any overt differences in the anatomical location or density of KLB-FLP neurons in males vs females (data not shown).

### GABAergic KLB neurons do not mediate response to protein restriction

To alternatively test the role of FGF21 signaling in GABAergic neurons, *Vgat-Cre* was crossed into the *Klb^lox/lox^* background to create *Klb^Vgat^* mice. Consistent with previous work, the LP diet reduced BW and lean mass gain, increased food intake, increased preference for casein, and increased EE in *Klb^lox/lox^* littermate controls (P < 0.05; **Figure 5**). These effects were also fully intact in *Klb^Vgat^* mice, with LP similarly altering body weight, food intake and preference, and energy expenditure in *Klb^lox/lox^* and *Klb^Vgat^* mice (**Figure 5**). These data indicate that FGF21 signaling (*Klb* expression) in GABAergic neurons is not required to respond to protein restriction.

**Figure 5.**
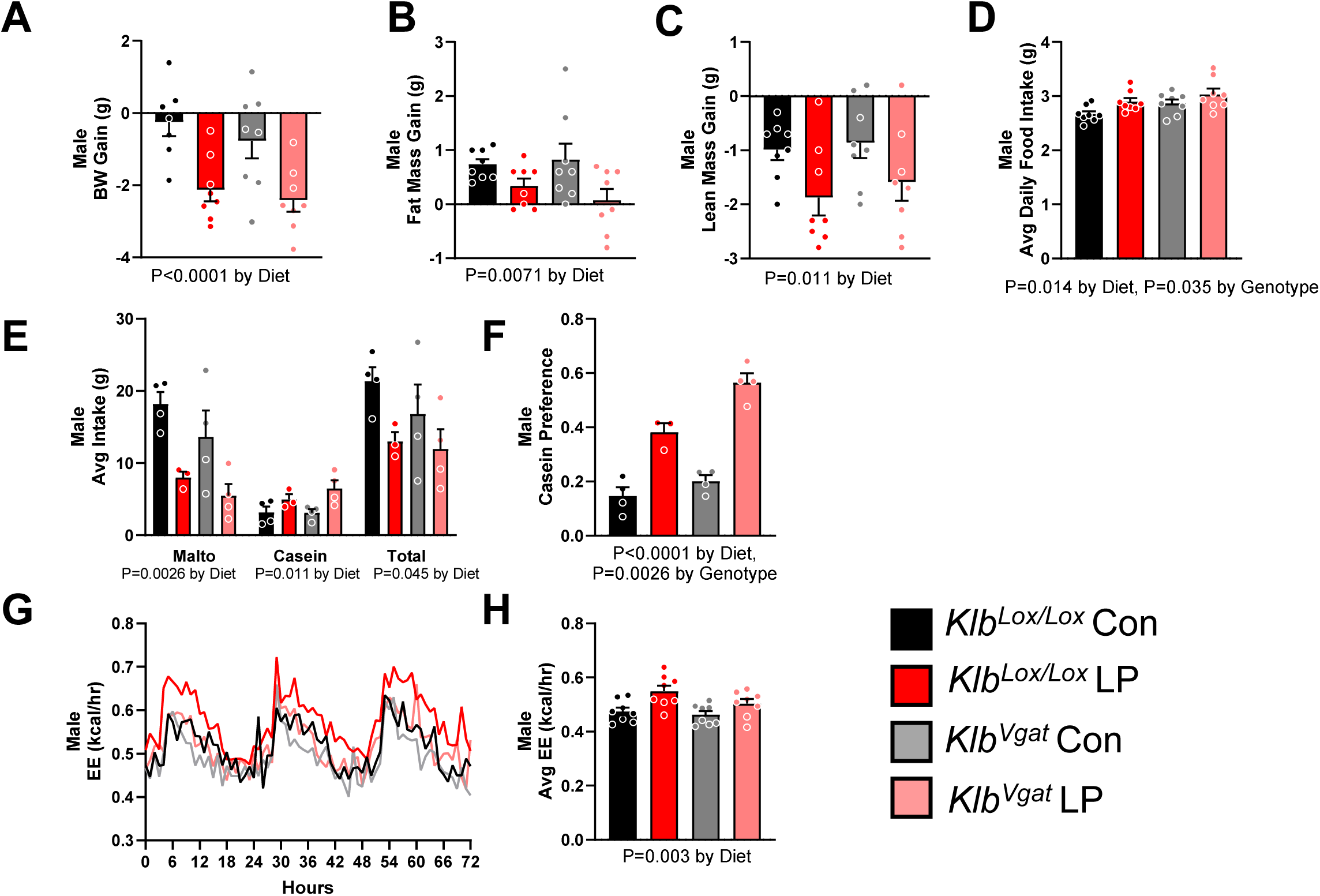
GABAergic KLB neurons do not mediate responses to low-protein diet. Male mice with *Vgat-Cre* specific deletion of *Klb* (*Klb^Vga^*^t^) had no difference in response to low protein diet (LP). Both *Klb^Vga^*^t^ and *Klb^lox/lox^* littermates lost a similar amount of weight when placed of LP diet (A) and this decrease was made of both a loss of fat mass (B) and lean mass (C). *Klb^Vga^*^t^ animals also responded like control animals to LP diet in their food intake by increasing overall food intake (D) and demonstrating a preference for casein (E & F) with overall effects of diet and genotype, but no interaction between the two. Finally, LP diet similarly elevated average energy expenditure (EE) in both groups over a 72-hour period (G & H). All data are mean ± SEM, analyzed with two-way ANOVA and post-hoc T-test, Avg EE was analyzed by ANCOVA and post-hoc T-test. n=4-8/group

### *Klb* expression is the VMH or PVN is not required for the metabolic response to protein restriction

Previous work has suggested that *Klb*-expressing neurons in both the ventromedial hypothalamus (VMH) and paraventricular nucleus (PVN) contribute to the effects of FGF21 on sweet/carbohydrate intake ^14, 15^. Although we did not detect *Klb-Flp*-induced recombination in these brain areas, we nevertheless tested whether FGF21 signaling in either the VMH or PVN was required for response to protein restriction. The VMH was targeted by crossing *SF1*-Cre ^26^ into the *Klb^lox/lox^* background, generating *Klb^Sf1-Cre^* mice. *Klb^Sf1-Cre^* and *Klb^lox/lox^* littermate controls were placed on Con or LP diet as above and fed for 6 weeks. LP diet again altered growth, food intake, and energy expenditure in male *Klb^lox/lox^* mice, and these effects were intact in *Klb^Sf1-Cre^* mice (P < 0.05, **Figure 6**). In female *Klb^lox/lox^*, LP increased food intake, and this effect was also intact in female *Klb^Sf1-Cre^* mice (**Figure S4 E & F**). These data indicate that *Klb* expression within the VMH is unnecessary for the adaptive response to protein restriction.

**Figure 6.**
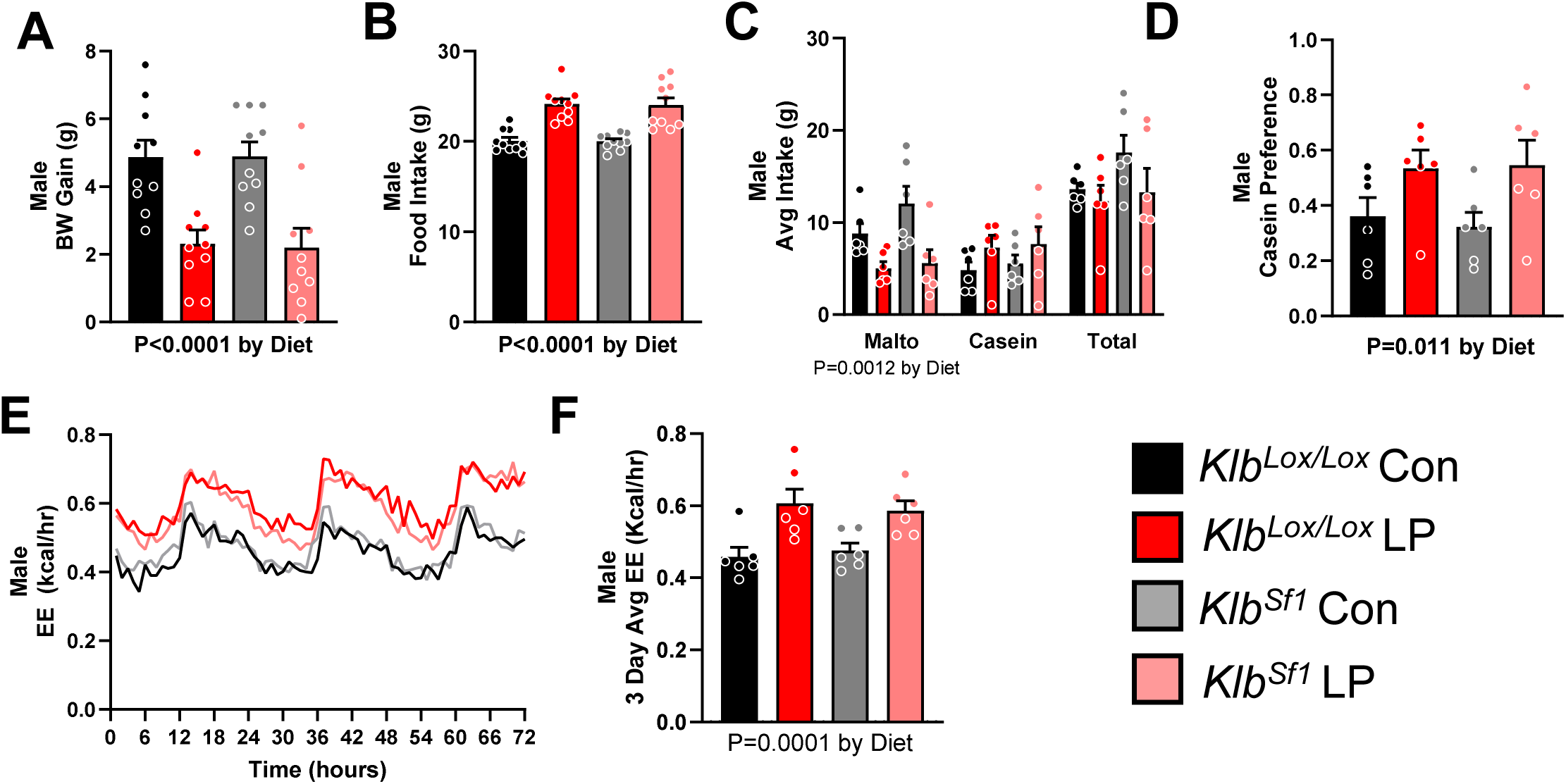
FGF21 signaling in the VMH does not mediate the effects of protein restriction. A VMH-specific KLB knockout was created by crossing *SF1-Cre* mice with *Klb*-floxed mice *(Klb^SF1^)*. When placed on LP diet, *Klb^SF1^* mice responded similarly to floxed controls in terms of body weight gain (A), food intake (B), protein preference (C & D), and energy expenditure (E & F). All data are mean ± SEM, analyzed with two-way ANOVA and post-hoc T-test, Avg EE was analyzed by ANCOVA and post-hoc T-test. n=6-10/group.

A similar experiment targeted the PVN using *Sim1-Cre* ^27^. Again, the established effects of LP diet to reduce BW gain but increase food intake, protein preference, and energy expenditure were observed in both *Klb^lox/lox^* and *Klb^Sim1-Cre^* mice (P < 0.05, **Figure 7**). However, the effects of LP diet to increase protein preference were slightly blunted in *Klb^Sim1-Cre^* mice compared to *Klb^lox/lox^* littermates (P < 0.01, **Figure 7 D)**. Taken together, these data indicate that *Klb* expression within the PVN is not required for mice to exhibit the adaptive response to protein restriction, with the exception of a modest effect on food choice.

**Figure 7.**
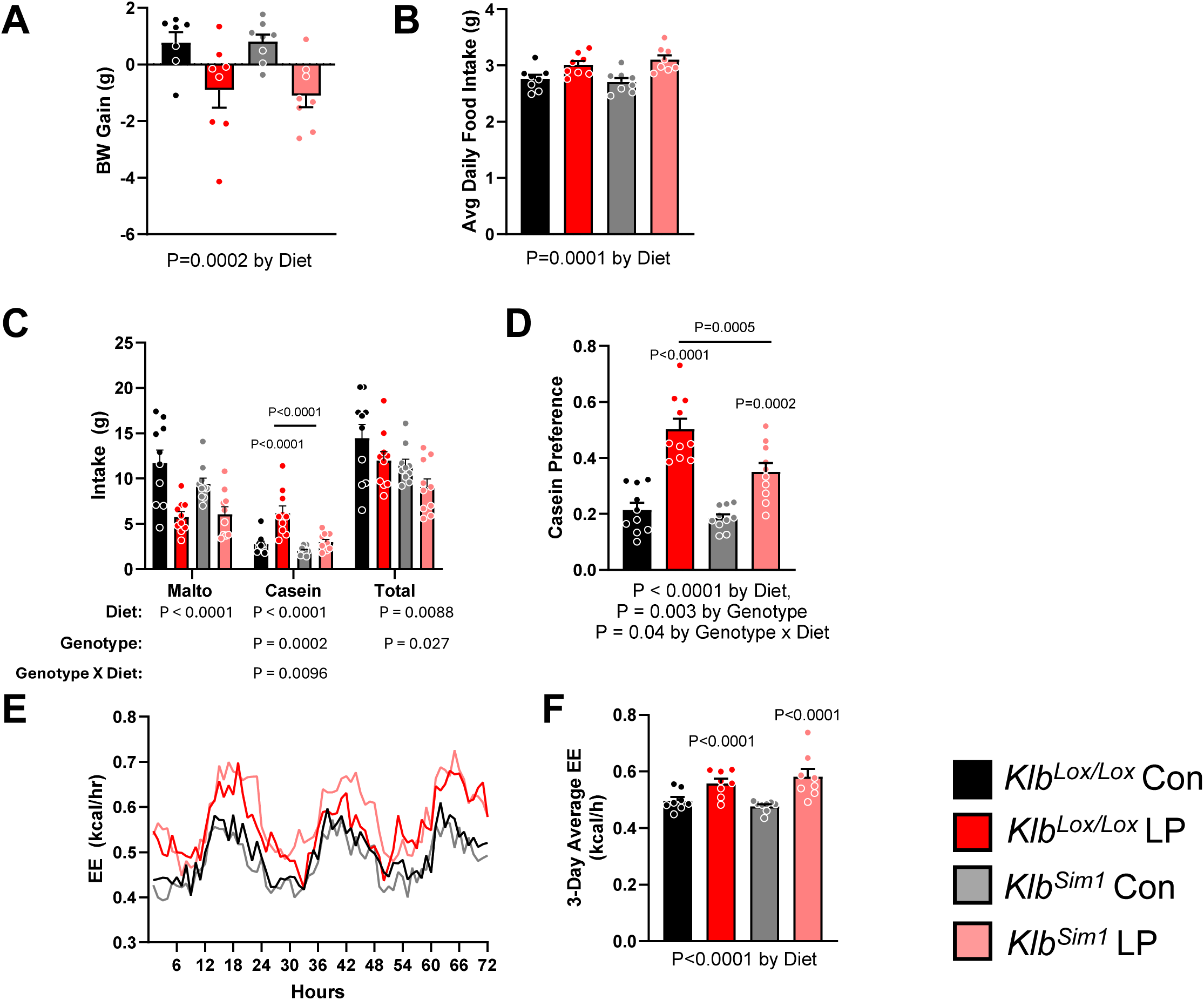
FGF21 signaling in the PVN does not mediate the effects of protein restriction. *Sim1-Cre* mice were crossed with *Klb*-floxed mice, and the resulting *Klb^Sim1^* were placed on LP diet. *Klb^Sim1^*mice responded similarly to floxed controls in terms of body weight gain (A), food intake (B), and EE (E & F). *Klb^Sim1^* mice did exhibit a more modest increase in casein consumption (C), reducing casein preference compared to *Klb^lox/lox^* controls on LP diet; however, the preference was still greater than in the control diet group (D). All data are mean ± SEM, analyzed with two-way ANOVA and post-hoc T-test, Avg EE was analyzed by ANCOVA and post-hoc T-test. n=8-10/group.

### Ablation of KLB neurons in the NTS blocks the effect of LP diet on food intake and EE

The above data provide little support for a model in which FGF21 signaling in the SCN, VMH, or PVN contributes to the response to protein restriction. Conversely, the NTS contains a large population of Klb neurons that are acutely responsive to FGF21 and marked by *Camk2a-Cre* and *Vglut2-Cre*. We, therefore, targeted these neurons for selective ablation via NTS-specific injections of a Flp-dependent AAV-Caspase3 or AAV-mCherry (control). LP diet reduced body weight gain in both AAV-Casp3 and AAV-mCherry injected mice (P = 0.028, **Figure 8 B**), suggesting that *Klb-Flp* neurons in the NTS are not required for effects of body weight gain. Conversely, the effects of LP diet on food intake (P < 0.01, **Figure 8 D**) and EE (P < 0.01, **Figure 8 E&F**) were both lost in AAV-Casp3 mice. There was also no difference in protein preference between AAV-Casp3 animals on Con and LP diet (**Figure 8 G&H**), however, this seems to be due to an increase in casein consumption in the mice on Con diet not lower consumption in the LP group. To confirm that AAV-Casp3 was not producing off-target effects, AAV-Casp3 was injected into the NTS of Flp-negative mice, where it had no effect on LP-induced increases in food intake and energy expenditure (**Figure S5**). Together, these data indicate that Klb-expressing neurons in the NTS are necessary for changes in food intake and energy expenditure in response to an LP diet.

**Figure 8.**
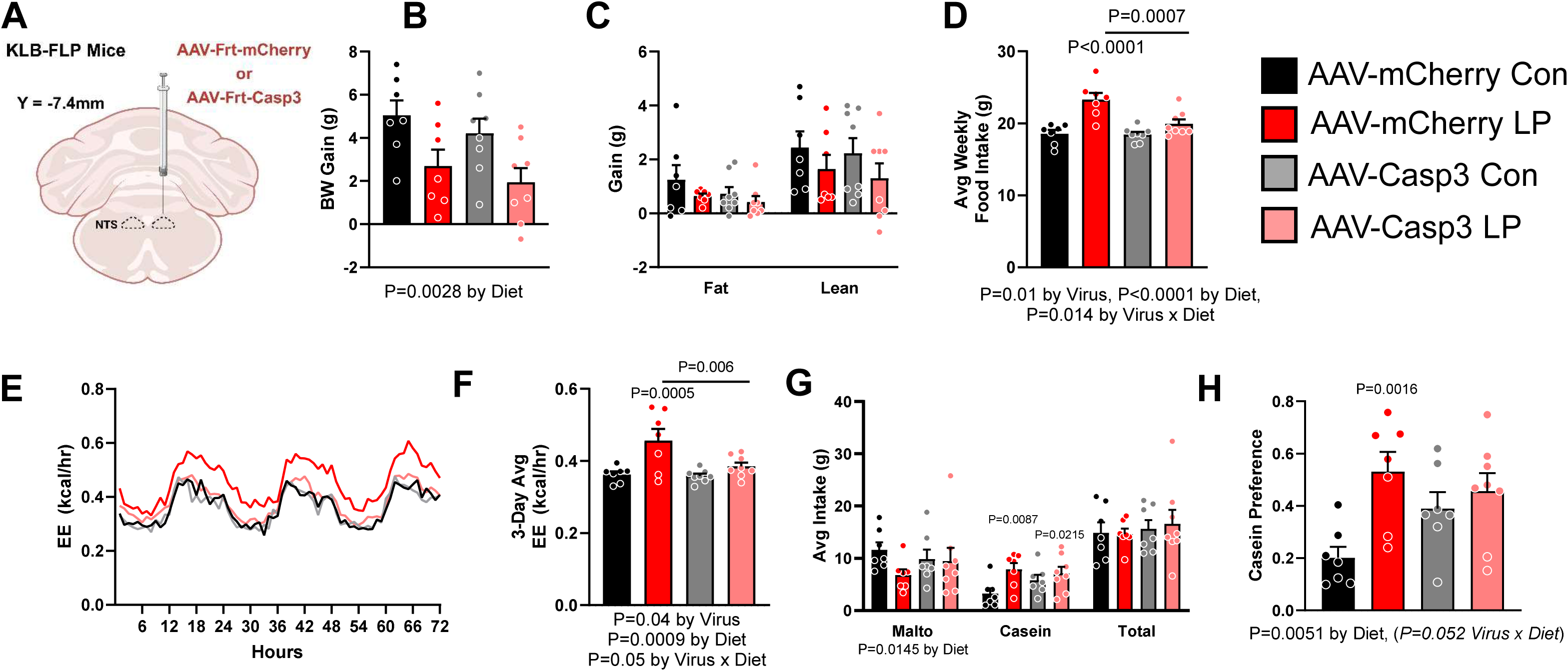
AAV-Casp3 ablation of NTS-KLB neurons blocks the effect of LP on food intake and EE. To target brainstem KLB neurons for ablation, Flp-dependent AAV-Casp3 or AAV-mCherry viruses were delivered to the NTS of *Klb-Flp* mice (A). Ablation of KLB neurons had no effect on body weight (B) or composition (C) in mice on LP diet. Mice that received the AAV-Casp3 virus lost the LP-induced increase in food intake (D) and EE (E&F) compared to the AAV-mCherry control group. All data are mean ± SEM, analyzed with two-way ANOVA and post-hoc T-test, Avg EE was analyzed by ANCOVA and post-hoc T-test. n=8/group.

### Chemogenetic activation of NTS KLB neurons increases EE and food intake

Since the above data suggest that NTS-KLB neurons are necessary for LP-induced changes in FI and EE, we next tested whether acute activation of these neurons is sufficient to drive changes in FI and EE. This question was first addressed by NTS-specific injection of a Flp-dependent AAV-hM3Dq virus or AAV-mCherry control in *Klb-Flp* mice. CNO-induced activation of neural activity was first confirmed via ex vivo electrophysiology three weeks following NTS-specific injection of AAV-hM3Dq, with CNO significantly increasing both firing frequency and resting membrane potential in *Klb-Flp* neurons (P < 0.05; **Figure S6 A-D**). Similarly, IP CNO injection significantly increased cFos expression in NTS-*Klb-Flp* neurons in mice receiving AAV-hM3Dq (P < 0.02; **Figure S6 E-H**).

AAV-hM3Dq or AAV-mCherry injected *Klb-Flp* mice were acclimated to Promethion metabolic chambers, treated with CNO injection (2, 3, 5 mg/kg IP), and EE assessed over 6 hours. CNO treatment increased EE in AAV-hM3Dq vs. AAV-mCherry mice across all doses (P = 0.012, **Figure 9 B**), having the most robust effect at the highest dose (P=0.0036, **Figure 9 B**). Conversely, CNO did not significantly alter food intake (**Figure 9 C**) during this 6hr period.

**Figure 9.**
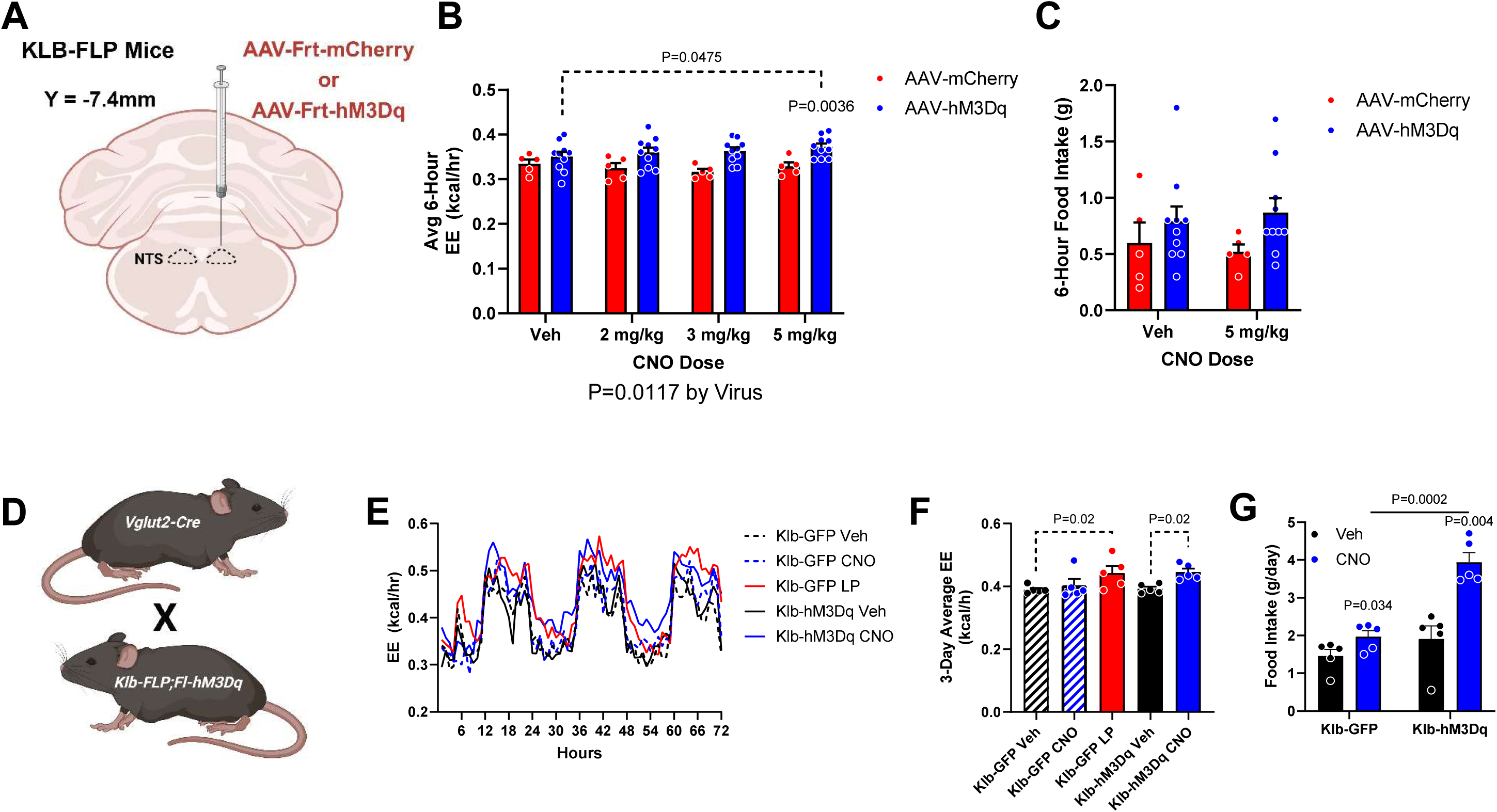
Chemogenetic activation of NTS-KLB neurons increases EE and food intake. Two methods were used to chemogenetically stimulate NTS-KLB neurons. In the first, Flp-dependent hM3Dq or mCherry viruses were delivered to the NTS of *Klb-Flp* mice (A), and acute i.p. administration of CNO (2, 3, 5 mg/kg) was tested in a crossover design. EE was elevated in the 6-hour period after the CNO injection with an overall effect of the virus and a significant difference at the highest dose when compared to mCherry controls (B). Acute administration of CNO did not effect food intake in the 6-hour period (C). In a second paradigm, NTS-KLB neurons were targeted for hM3Dq expression via a 3-way cross of *Vglut2-Cre;Klb-Flp;FL-hM3Dq* (D), with CNO delivered chronically in the drinking water (11.25 ug/ml). EE was elevated in *Vglut2-Cre;Klb-Flp;FL-hM3Dq (Klb-hM3Dq)* mice treated with CNO compared to those given untreated drinking water or *Klb-Flp;FL-hM3Dq (Klb-GFP)* mice lacking Vglut-Cre, and this increased EE was to a similar degree as littermate controls fed LP diet (F). Chronic CNO also increased the average daily food intake in *Vglut2-Cre;Klb-Flp;FL-hM3Dq (Klb-hM3Dq)* animals (G). All data are mean ± SEM, analyzed with paired two-way ANOVA and post-hoc T-test. Avg EE was analyzed by ANCOVA and post-hoc T-test. n= 5-10/group.

A second approach utilized intersectional genetics to target NTS-KLB neurons via the dual recombinase RC::FL-hM3Dq line ^28^, in which Flp-induced recombination drives GFP expression but the addition of Cre removes the GFP and induces an hM3Dq-mCherry fusion protein. Since *Klb-Flp* neurons in the NTS are the only neural population labeled by *Klb-Flp* and *Vglut2-Cre*, a triple cross of *Klb-Flp*;*Vglut2-Cre*;FL-hM3Dq was generated to target hM3Dq expression to this NTS population (**Figure S6 H**). Male *Klb-Flp*;*Vglut2-Cre*;FL-hM3Dq mice and Cre-negative littermate controls (*Klb-Flp*;FL-hM3Dq) were placed into metabolic cages at 8 weeks of age and provided CNO in the drinking water (11.25 ug/ml). While CNO treatment did not affect EE in Cre-control mice, CNO exposure significantly increased EE in *Klb-Flp*;*Vglut2-Cre*;FL-hM3Dq mice over the 3-day experiment (P =0.02; **Figure 9 E&F**), and this effect was equivalent in size to that induced by LP diet (**Figure 9 F**). CNO also significantly increased average 24hr food intake in *Klb-Flp*;*Vglut2-Cre*;FL-hM3Dq mice but not in Cre-littermates (P < 0.05, **Figure 9 G**). Thus, two separate experiments suggest that chemogenetic activation of NTS-KLB neurons is sufficient to drive metabolic endpoints in a manner analogous to LP diet.

## Discussion

The ability to detect and adapt to changes in nutrient availability is fundamental to survival. While the mechanisms underlying the detection of energy, water, and salt restriction are well-characterized, our understanding of how animals detect and respond to protein restriction has lagged behind, despite the critical importance of protein in human health and nutrition. Our previous work identified FGF21 as an essential mediator of the adaptive response to protein restriction, with protein-restricted mice exhibiting robust increases in circulating FGF21 that drive changes in both food intake and energy expenditure ^1, 2, 3^. Whole body deletion of FGF21, or the brain-specific deletion of its co-receptor Klb, completely blocks the adaptive response to dietary protein restriction in mice. However, the specific neural circuits through which FGF21 coordinates these responses remained unclear. Here we identify a discrete population of glutamatergic, Klb-expressing neurons in the nucleus of the solitary tract (NTS) that are both necessary and sufficient for key metabolic adaptations to protein restriction.

To identify the neural circuits mediating FGF21 action, we developed a novel *Klb-Flp* mouse line that allows precise targeting of FGF21-responsive neurons through intersectional genetic approaches. This approach revealed two primary populations of Klb-expressing neurons: one in the suprachiasmatic nucleus (SCN) and another in the NTS. Importantly, these populations are phenotypically distinct, with NTS neurons being glutamatergic while SCN neurons are GABAergic. This distinction proved crucial, as deletion of *Klb* from glutamatergic but not GABAergic neurons blocked the adaptive response to protein restriction, consistent with previous work ^22^. The selective loss of protein-restriction responses in mice lacking *Klb* in glutamatergic neurons, combined with our observation that *Klb*-expressing neurons in the NTS are exclusively glutamatergic, strongly implicated the NTS as a key site of FGF21 action.

Several complementary lines of evidence further strengthened this conclusion. First, FGF21 directly activated NTS-KLB neurons both in vivo (cFos) and ex vivo (electrophysiology). This effect is likely a direct action on Klb neurons, since the effects on firing rate and membrane potential persisted even when synaptic transmission was blocked. However, these data do not exclude the additional possibility of indirect effects from other neural populations or FGF21-sensitive vagal/DRG afferents. Interestingly, we did not observe clear effects of FGF21 within the SCN, as there was no effect on cFos and mixed effects on neural activity during electrophysiological analysis. Second, selective ablation of NTS-KLB neurons using a Flp-dependent caspase prevented protein restriction-induced changes in food intake and energy expenditure. Finally, chemogenetic activation of these neurons was sufficient to increase both food intake and energy expenditure, mimicking the key metabolic effects of protein restriction. Together, these data provide compelling evidence that FGF21 acts through Klb-expressing neurons in the NTS to coordinate changes in food intake and energy expenditure in response to protein restriction.

Our findings challenge previous assumptions about the brain areas mediating FGF21 action. Early work using transgenic FGF21 overexpression suggested that the NTS was not critical for FGF21’s effects ^10^, leading to a focus on other brain regions including the SCN, PVN, and VMH ^10, 12, 14, 15, 16, 17, 18^. However, our work using the *Klb-Flp* reporter line, direct manipulation of neural activity, and genetic deletion of *Klb* provide little support for a model in which FGF21 signaling in these areas plays a key role in the response to protein restriction. Indeed, the deletion of *Klb* from either the VMH (using *SF1-Cre*) or PVN (using *Sim1-Cre*) did not affect protein restriction responses. Notably, none of these prior studies focused on a model of dietary protein restriction, which produces a sustained, physiological increase in circulating FGF21 levels. While we cannot rule out contributions from other brain regions, the lack of effect in these hypothalamic areas led to our reevaluation of the NTS, and our data establish the NTS as a critical site for FGF21 action during protein restriction.

Identifying the NTS as a key mediator of FGF21 action is particularly intriguing given this region’s established role in nutrient sensing and metabolism. The NTS receives direct vagal input from the gastrointestinal tract and serves as a primary integration center for visceral signals related to feeding and metabolism. While additional work is needed to identify the neural inputs and outputs of NTS-KLB neurons, our finding that FGF21 acts through glutamatergic neurons in this region adds to growing evidence that the NTS contains distinct neural populations that coordinate specific aspects of nutrient homeostasis. Interestingly, while these NTS-KLB neurons express Camk2a, they were not effectively labeled by *Phox2b-Cre* despite prior evidence that *Phox2b-Cre* reduces *Klb* expression within the NTS ^10^. Additional work is required to more deeply phenotype this population of Camk2a/Vglut2-positive Klb neurons, as these neurons may represent a unique population within the NTS specifically tuned to protein status.

An unexpected finding in our study was the dissociation between growth effects and metabolic effects following the ablation of NTS-KLB neurons. While caspase-mediated ablation of these neurons blocked protein restriction-induced changes in food intake and energy expenditure, it did not prevent the reduction in growth typically seen with protein restriction. This contrasts with work using *Camk2a-Cre* or *Vglut2-Cre*, where all protein restriction responses were blocked ^2^. Several possibilities could explain this dissociation. First, *Camk2a-Cre* or *Vglut2-Cre* represent germline deletions present from birth, whereas AAV-Caspase injection ablates neurons in adulthood. Second, brain microinjections may not hit all neurons within a brain area, particularly with a structure such as the NTS. This lack of coverage may explain the lack of effect on some endpoints and the generally weaker effects of microinjection vs. germline manipulation. Finally, the growth response to protein restriction may be mediated by a distinct population of FGF21-responsive neurons not marked by our *Klb-Flp* line. This possibility is particularly intriguing given that our *Klb-Flp* line appears to mark a much narrower set of neurons than previously published *Klb-Cre* lines ^14^. While this suggests our *Klb-Flp* mice may miss populations with lower expression levels, this line provides high specificity for Klb neurons in the NTS and SCN, which are the two areas most clearly marked via *Klb* in situ hybridization ^9, 10^.

Notably, our findings primarily focus on males, where protein restriction produces robust effects on multiple metabolic endpoints. Prior work indicates that females exhibit a more modest response to protein restriction, particularly in terms of metabolic endpoints ^23, 24^, but that effects of protein restriction on food intake and food choice remain ^25^. Our work is largely consistent with this evidence, as females mice did not exhibit changes in body weight gain on LP, but did exhibit a hyperphagic response. Importantly, this hyperphagic response nevertheless requires *Klb* expression in NTS neurons and *Vglut2-Cre* neurons. This observation suggests that while the magnitude of protein restriction responses may differ between sexes, the fundamental neural circuit mediating these responses remains consistent.

Our findings also raise important questions about the broader neural circuits through which NTS-KLB neurons influence metabolism and behavior. These neurons likely engage downstream circuits controlling both food intake and energy expenditure. Indeed, the fact that both food intake and energy expenditure increase following chemogenetic activation of these neurons suggests they may coordinate multiple parallel circuits to achieve metabolic homeostasis during protein restriction. Future studies mapping the precise connectivity of these neurons and determining how they integrate with other nutrient-sensing circuits will be crucial for understanding the full scope of their function.

The ability to detect and respond to changes in nutritional state is a core feature of biology, allowing animals to match behavior and metabolism to ongoing nutrient availability ^21, 29^. Our work demonstrates that a discrete population of *Klb*-expressing NTS neurons are responsive to FGF21 and coordinate changes in food intake and energy expenditure in response to protein restriction. By identifying this critical neural circuit, we begin to uncover how protein status is decoded by the brain, advancing our understanding of a fundamental biological process essential for survival.

## Methods

### Animals

All animal studies were approved by the PBRC Institutional Animal Care and Use Committee and were performed following the guidelines and regulations of the NIH Office of Laboratory Animal Welfare. Male and female mice were 8-12 weeks of age and maintained on a standard chow diet (Purina 5001) until transfer to experimental diets. All mice were bred on a C57BL/6 background. *Klb-Flp* mice were generated by inserting Flp recombinases into the 3’ end of the Klb gene, separated by a T2A linker, resulting in the expression of Flp in all cells expressing Klb. *Klb-Flp* mice were also crossed with two different dual recombinase lines. RC::FLTG mice express tdTomato in response to FLP but GFP in response to Flp/Cre coexpression ^19^. RC::FL-hM3Dq express GFP in response to Flp recombination but hM3Dq-mCherry in response to combined Flp/Cre recombination ^28^. Additional Cre lines (*Vglut2-Cre*, *Vgat-Cre*, *Camk2a-Cre*, and *Phox2b-Cre*) were purchased from Jackson Labs and used to create intersectional crosses and phenotype Klb neurons. Klb^lox/lox^ mice were provided by Dr. Steven Kliewer ^7, 10^ and crossed with various Cre lines (*Vglut2-Cre*, *Vgat-Cre*, *Sim1-Cre*, *SF1-Cre*) to generate tissue-specific Klb knockouts. Animals were single-housed in a 12:12hr light:dark cycle with ad libitum access to food and water unless otherwise noted.

### Experimental Diets

Diets were formulated and produced by Research Diets and were designed to be isocaloric by adjusting protein and carbohydrate content while keeping fat constant. The Control diet contained 20% casein (by weight) as the protein source, while the low protein (LP) diet contained 5% casein. Diet compositions have been previously published ^2, 4, 5^.

### Stereotaxic Surgery

Mice were anesthetized with vaporized isoflurane and positioned in the stereotaxic frame. The skull was exposed and cleaned with 0.03% H2O2. A hole was drilled through the skull at coordinates determined relative to Bregma. For viral injections targeting the NTS, the coordinates were: AP -7.4 mm, ML ±0.3 mm, and DV - 4.75 mm from the surface of the brain. A 1 μL Hamilton syringe connected to an injector was used to deliver 300 nL of virus per injection site. The following viral constructs were used:

- AAV-Casp3 (AAV-8/2-hEF1α-dFRT-(pro)taCasp3_2A_TEVp(rev)-dFRT-WPRE-hGHp(A))
- AAV-mCherry control (AAV-8/2-hSyn1-dFRT-mCherry(rev)-dFRT-WPRE-hGHp(A))
- AAV-hM3Dq (AAV-8/2-hSyn1-dFRT-hM3D(Gq)_mCherry(rev)-dFRT-WPRE-hGHp(A))

For ablation studies, *Klb-Flp* mice received NTS-specific injections of AAV-Casp3 or AAV-mCherry. For chemogenetic activation studies, mice received either Flp-dependent AAV-hM3Dq or AAV-mCherry control. All viruses were provided by the the Viral Vector Facility of the Neuroscience Center Zurich (ZNZ), University of Zurich.

### Indirect Calorimetry

Energy expenditure was measured using Promethion metabolic chambers (Sable Systems). Mice were acclimated to the chambers for at least 3 days before measurements. For chemogenetic activation studies, CNO was administered either by IP injection (2-5 mg/kg) or in drinking water (11.25 ug/ml). Data were analyzed using the CalR for initial quality control and mean generation, with EE being subsequently analyzed via ANCOVA with body weight as a covariate.

### Nutrient Preference Tests

For two-bottle choice tests, mice were simultaneously offered solutions containing either 4% casein/0.1% saccharin or 4% maltodextrin/0.1% saccharin for 3 days. Bottle positions were counterbalanced and swapped daily. Consumption was measured daily and averaged across the test period. Preference ratios were calculated as (casein consumption)/(total consumption).

### Electrophysiology

Acute brain slices (300 μm) containing the NTS or SCN were prepared from Klb-Flp;FLTG mice. Briefly, mice were anesthetized by isoflurane and transcranial perfused with a modified ice-cold sucrose-based cutting solution (pH 7.3) containing 10 mM NaCl, 25 mM NaHCO3, 195 mM Sucrose, 5 mM Glucose, 2.5 mM KCl, 1.25 mM NaH2PO4, 2 mM Na-Pyruvate, 0.5 mM CaCl2, and 7 mM MgCl2, bubbled continuously with 95% O2 and 5% CO2 ^30^. The mice were then decapitated, and the entire brain was removed and immediately submerged in the cutting solution. Slices (250 µm) were cut with a Micron Leica VT1000s vibratome (Leica, IL, U.S.). Brain slices containing the DRN were obtained for each animal (Bregma -2.06 mm to -1.46 mm; Interaural 1.74 mm to 2.34 mm). The slices were recovered for 1 h at 34°C and then maintained at room temperature in artificial cerebrospinal fluid (aCSF, pH 7.3) containing 126 mM NaCl, 2.5 mM KCl, 2.4 mM CaCl2, 1.2 mM NaH2PO4, 1.2 mM MgCl2, 5.0 mM glucose, and 21.4 mM NaHCO3) saturated with 95% O2 and 5% CO2 before recording. Whole-cell patch-clamp recordings were performed at 32-34°C in artificial cerebrospinal fluid (aCSF). The current-clamp mode was engaged to measure neural firing frequency and resting membrane potential. The values for RM and firing frequency averaged within the 2-min bin. FGF21 (300 nM) was puff delivered and changes in membrane potential and firing frequency were recorded. For synaptic blockade experiments, CNQX (10 μM), AP5 (50 μM), and picrotoxin (100 μM) were added to the aCSF.

### Immunohistochemistry

Mice were transcardially perfused with 10% formalin and brains were post-fixed overnight. Following cryoprotection in 30% sucrose, brains were sectioned at 30 μm on a sliding microtome. For cFos studies, mice received IP injection of either FGF21 (1 mg/kg) or saline 2 hours before perfusion. Sections were immunostained with antibodies against cFos (1:1000, Synaptic Systems), dsRed (1:1000, Takara), and GFP (1:1000, Abcam). Secondary antibodies conjugated to biotin or fluorophores were used at 1:200 dilution.

### RNAscope In Situ Hybridization

RNAscope HiPlex v2 reagents were used to colocalize *Klb* mRNA with tdTomato expression in *Klb-Flp;FLTG* mice following manufacturer’s protocols (Advanced Cell Diagnostics). Sections were imaged using a Leica TIRF/DM6000 microscope.

### Statistical Analysis

Data were analyzed using the SAS software package (SAS V9, SAS Institute) using one-way, two-way, or repeated measures ANOVA using the general linear model procedure. When experiment-wide tests were significant, post-hoc comparisons were made using the LSMEANS statement with the PDIFF option, and represent least significant differences tests for pre-planned comparisons. Average daily energy expenditure was analyzed via analysis of covariance (ANCOVA) with body weight as the covariate using the general linear model procedure of SAS. All data are expressed as mean ± SEM, with a probability value of 0.05 considered statistically significant. Groups sizes are described in their respective figure legend.

## Acknowledgments

The authors would like to thank the leadership and staff of the PBRC Comparative Biology Core and Animal Metabolism and Behavior Core for their skillful assistance and excellent technical support. This work was supported by the National Institutes of Health (NIH) R01DK123083, R01DK121370, and S10OD023703 to CDM. RAS was supported by F32DK130544. SQK was supported by T32DK064584. DHM was supported by R01DK131165. HM was supported by R01AT011683 and R01DK092587. YH was supported by R01DK129548. This project used facilities within the Animal Metabolism & Behavior Core, Genomics Core, and Cell Biology and Bioimaging Core at PBRC that are supported in part by NIH center awards P20GM135002 and P30DK072476, as well as an NIH equipment award S10OD023703.

## Author Contributions

Conceptualization – RAS, SY, CDM; Methodology – RAS, SQK, MSK, HRB, YH, SY, CDM; Investigation - RAS, SQK, MSK, DAA, YH, SY, CDM; Visualization – RAS, HRB, DM, YH, HM, SY, CDM; Formal - analysis – RAS, YH, SY, CDM; Writing - Original Draft – RAS, YH, SY, CDM; Writing - Review & Editing – RAS, SQK, MSK, DAA, HRB, DM, YH, HM, SY, CM. Funding acquisition – RAS, CDM.

## Supplemental Figures

**Supplemental figure 1.**
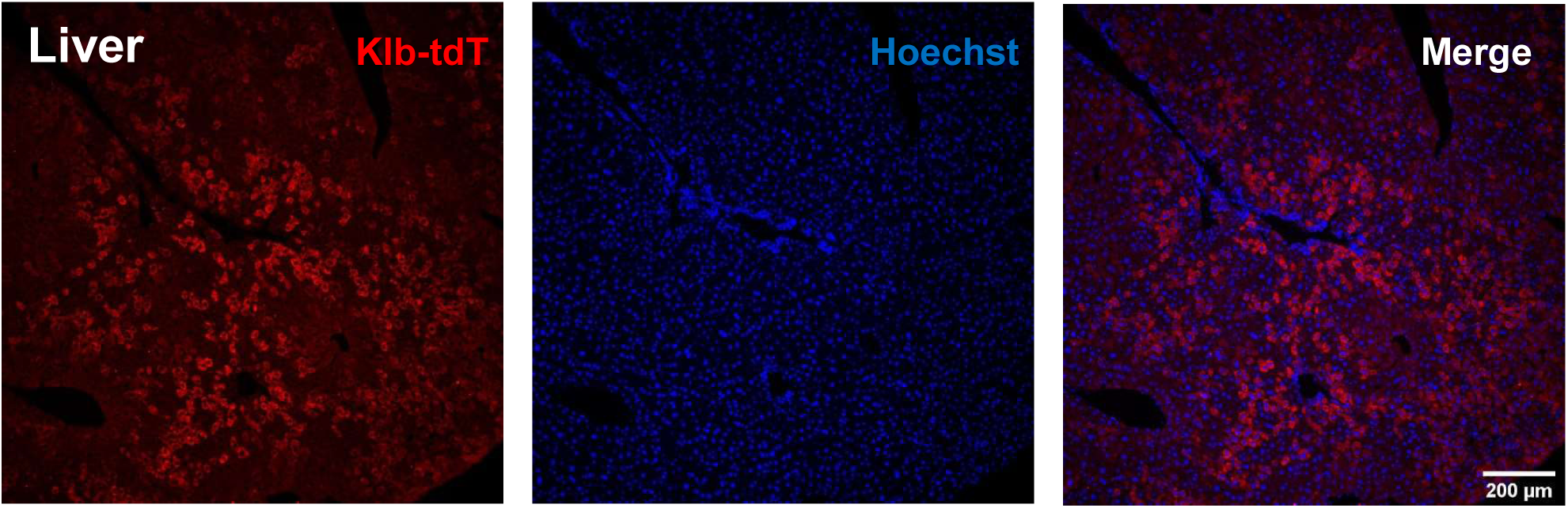
*Klb-Flp* reporter expression in the liver. Along with tdTomato expression in the brain, reporter expression was also detected throughout the liver.

**Supplemental Figure 2.**
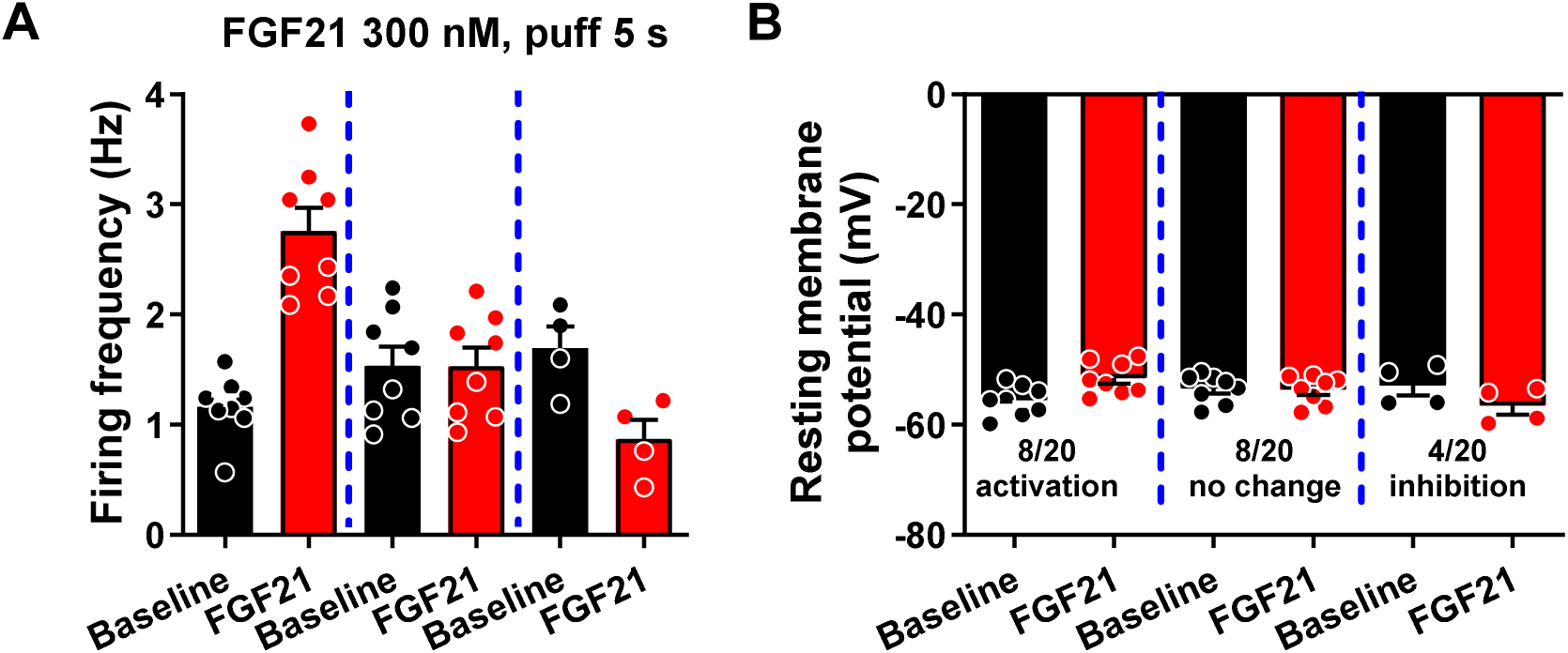
FGF21-dependent regulation Klb neurons in the SCN. Slice patch-clamp electrophysiology was performed in the SCN of *Klb-Flp;FLTG* mice with bath administration of hFGF21 (300nM). FGF21 produced divergent effects on both firing frequency (A) and resting membrane potential (B).

**Supplemental Figure 3.**
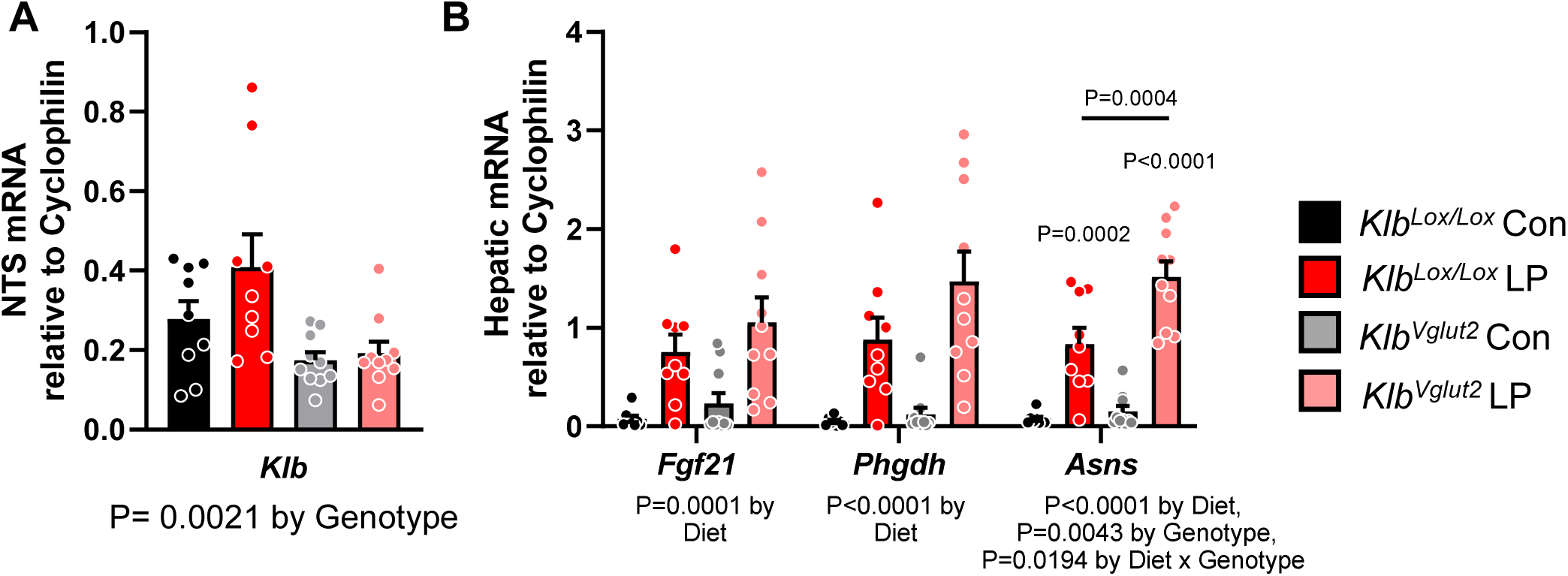
qPCR in *Vglut2*-Cre mice. RNA was extracted from the NTS and the liver of *Vglut2*-Cre and littermate control mice. We confirmed that there was a reduction in *Klb* expression in the NTS of Cre-positive animals (A). We next confirmed that animals were responding to protein deprivation with an increased expression of *Fgf21*, phosphoglycerate dehydrogenase (*Phgdh*), and asparagine synthetase (*Asns*) within the liver (B). All data are mean ± SEM, analyzed with two-way ANOVA and post-hoc T-test.

**Supplemental Figure 4.**
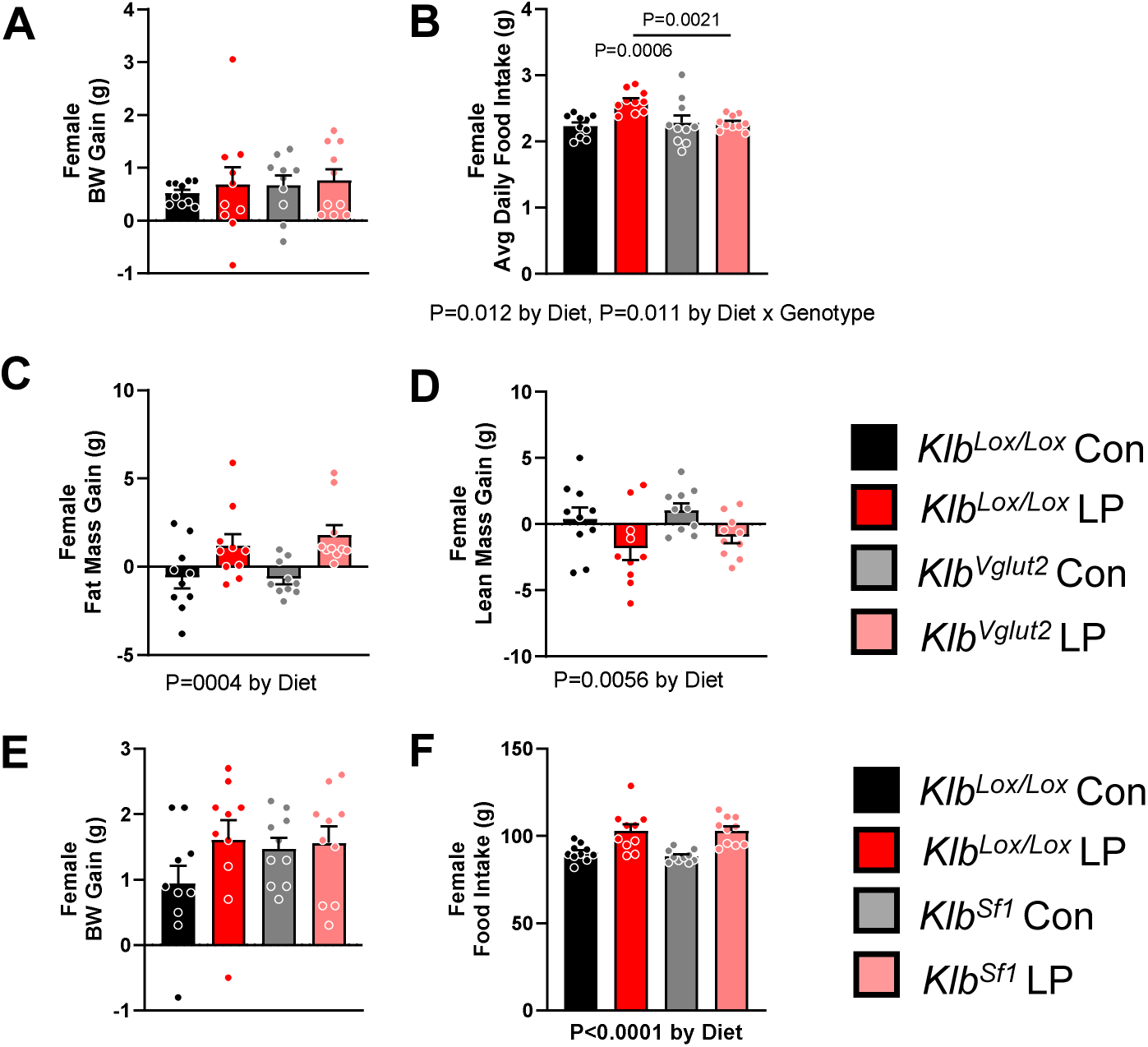
Effect of protein restriction in female *Klb^Vglut2-Cre^* and *Klb^Sf1-Cre^* mice. In female mice, LP diet had no effect on body weight gain, regardless of genotype (A, E). Unlike males, LP diet increased in fat mass (C) and a decreased in lean mass (D) in females, and this effect was is not altered in *Klb^Vglut2-Cre^* females. Like males, LP-fed females increased food intake, and this effect was lost in *Klb^Vglut2-Cre^* females (B), but not in *Klb^Sf1-Cre^* females (F). All data are mean ± SEM, analyzed with two-way ANOVA and post-hoc T-test.

**Supplemental Figure 5.**
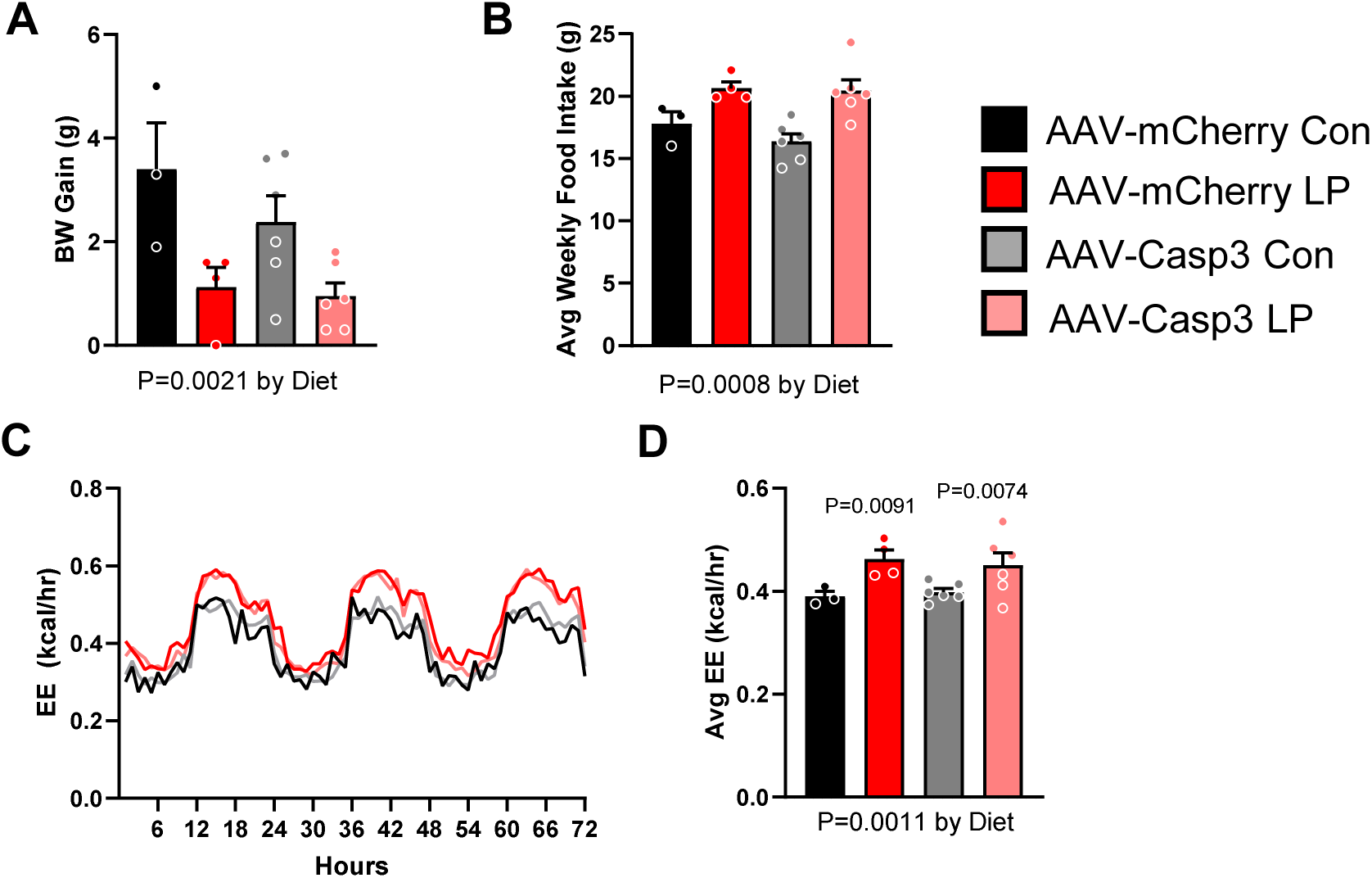
Control experiment for Flp-dependent AAV-Casp3. Wildtype C57Bl/6 mice were used as a control group to confirm that AAV-Casp3 had no off-target effects. Mice that received AAV-Casp3 virus responded like AAV-mCherry controls when placed on LP diet. In both groups, LP diet reduced BW gain (A) and increased food intake (B) and EE (C&D). All data are mean ± SEM, analyzed with two-way ANOVA and post-hoc T-test, Avg EE was analyzed by ANCOVA and post-hoc T-test.

**Supplemental Figure 6.**
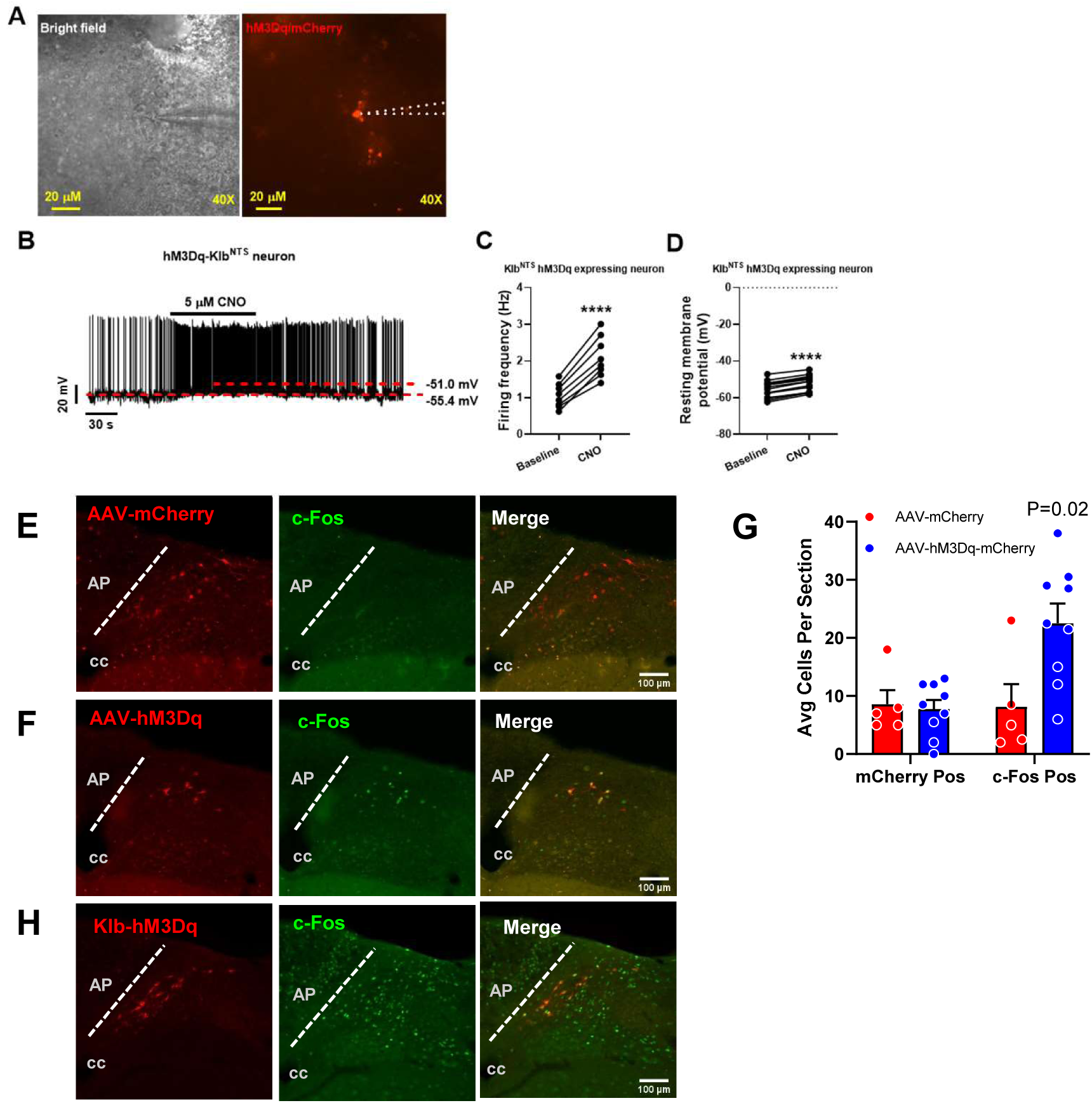
Validating chemogenetic activation of KLB neurons. To confirm chemogenetic stimulation of NTS-KLB neurons following AAV-hM3Dq injection, patch-clamp recording was performed on *Klb-Flp* animals following NTS-specific injection of AAV-hM3Dq (A). CNO increased firing frequency (B & C) and resting membrane potential (D). In vivo, CNO injection significantly increased NTS c-Fos expression in Klb-Flp mice injected with AAV-hM3Dq compared to AAV-mCherry mice (E, F, & G). Finally, CNO injection also increased c-Fos in the NTS of *Vglut2-Cre;Klb-Flp;FL-hM3Dq* animals (H).

